# DeepFoci: Deep Learning-Based Algorithm for Fast Automatic Analysis of DNA Double Strand Break Ionizing Radiation-Induced Foci

**DOI:** 10.1101/2020.10.07.321927

**Authors:** Tomas Vicar, Jaromir Gumulec, Radim Kolar, Olga Kopecna, Eva Pagáčová, Martin Falk

## Abstract

DNA double-strand breaks, marked by Ionizing Radiation-Induced (Repair) Foci (IRIF), are the most serious DNA lesions, dangerous to human health. IRIF quantification based on confocal microscopy represents the most sensitive and gold standard method in radiation biodosimetry and allows research of DSB induction and repair at the molecular and a single cell level. In this study, we introduce DeepFoci - a deep learning-based fully-automatic method for IRIF counting and its morphometric analysis. DeepFoci is designed to work with 3D multichannel data (trained for 53BP1 and γH2AX) and uses U-Net for the nucleus segmentation and IRIF detection, together with maximally stable extremal region-based IRIF segmentation.

The proposed method was trained and tested on challenging datasets consisting of mixtures of non-irradiated and irradiated cells of different types and IRIF characteristics - permanent cell lines (NHDF, U-87) and cell primary cultures prepared from tumors and adjacent normal tissues of head and neck cancer patients. The cells were dosed with 1-4 Gy gamma-rays and fixed at multiple (0-24 h) post-irradiation times. Upon all circumstances, DeepFoci was able to quantify the number of IRIF foci with the highest accuracy among current advanced algorithms. Moreover, while the detection error of DeepFoci remained comparable to the variability between two experienced experts, the software kept its sensitivity and fidelity across dramatically different IRIF counts per nucleus. In addition, information was extracted on IRIF 3D morphometric features and repair protein colocalization within IRIFs. This allowed multiparameter IRIF categorization, thereby refining the analysis of DSB repair processes and classification of patient tumors with a potential to identify specific cell subclones.

The developed software improves IRIF quantification for various practical applications (radiotherapy monitoring, biodosimetry, etc.) and opens the door to an advanced DSB focus analysis and, in turn, a better understanding of (radiation) DNA damaging and repair.

**Highlights:**

- New method for DSB repair focus (IRIF) detection and multi-parameter analysis
- Trainable deep learning-based method
- Fully automated analysis of multichannel 3D datasets
- Trained and tested on extremely challenging datasets (tumor primary cultures)
- Comparable to an expert analysis and superb to available methods

**Graphical Abstract:** 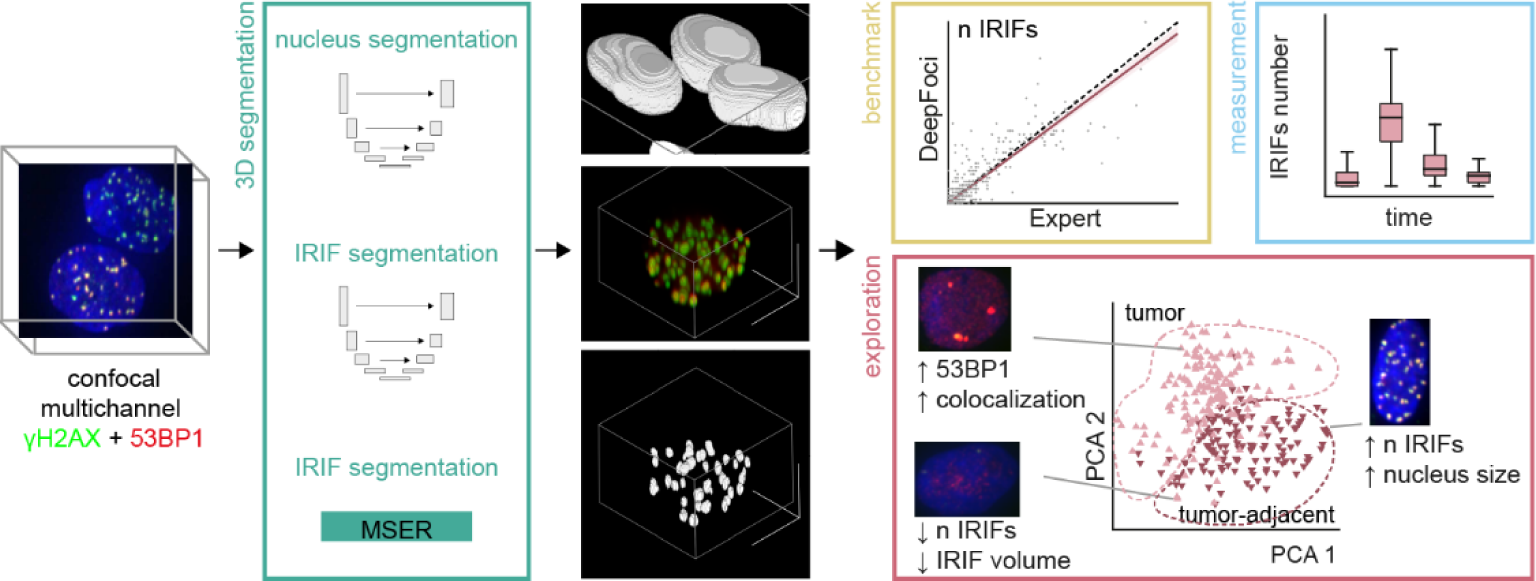

## Introduction

The ability to precisely and rapidly monitor DNA double-strand break (DSBs) induction and repair undermines numerous fields of biological, medical and space research (reviewed e.g., in [1–4]). DNA double-strand breaks (DSBs) are permanently introduced into the DNA molecule by ionizing radiation, radiomimetic chemicals and vital cell processes [5–7]. Simultaneously cutting both DNA strands, DSBs represent the most serious type of DNA lesions [8], which accumulation, if left unrepaired, fuels ageing [9], neurodegeneration [10], infertility [11], and other health consequences. Imprecise DSB repair then causes mutagenesis and may lead to cancer [7,12]. DSB damage induction and repair monitoring thus opens the doors to personalized medicine [3,4,13–15], for instance, coupled with direct radiotherapy effect monitoring [13,16], and rational development of new DNA damaging [17–20] or protecting drugs [21–24] needed in medicine [25,13,16] and radiation protection [26,27]. Precise and automated DSB damage monitoring is also irreplaceable in biodosimetry [26–30], for instance in situations related to mass radiation accidents (terrorist attacks with radioactive materials) or space exploration where astronauts would be exposed to a mixed field of different radiation types [31–37].

A revolution in DSB detection has come with the discovery of ionizing radiation-induced (repair) foci (IRIFs) that rapidly form at DSB sites after damage and currently serve as their most sensitive markers (reviewed, e.g., in [2–4]). One of the early events at DSB sites is the phosphorylation of histone H2AX at serine 139 (referred to as γH2AX) that eventually spreads over 2 Mb of damaged chromatin and leads to the formation of so-called γH2AX foci [38]. γH2AX foci then serve as signaling and structural platforms [39] that attract, in a spatially and temporally organized manner, additional repair proteins to DSB sites. Consequently, IRIFs of different repair proteins, characterized by specific parameters and behavior, can be visualized in cell nuclei by immunofluorescence microscopy. Since the number of IRIFs tightly corresponds with the number of DSBs in most DNA damage situations [40–42], the IRIFs formed by γH2AX or co-localized repair proteins (like 53BP1) can be considered quantitative DSB markers with a single lesion sensitivity [43].

While flow cytometry offers fast automated quantification of integrated values of these repair signals in high cell numbers [44], microscopy allows detection of individual IRIFs in the context of their natural chromatin environment in individual cells and analysis of their properties development in time [34,45,46]. Characterization of morphological and behavioral parameters of IRIFs formed by repair proteins participating in different DSB repair pathways [47]—such as 53BP1 and γH2AX, which were used in the present manuscript for illustration—opens the doors to the exploration of spatiotemporal interactions between repair proteins at individual DSB sites, deepening our insights into mechanisms of DSB induction and (mis)repair [13–15,36,48–55].

IRIFs are by nature highly dynamic structures. Their number per nucleus and principle parameters, such as the size, intensity, shape, and border sharpness change dramatically during post-damage time as repair proceeds [36,56–59]. Moreover, cell types or even individual cells of the same population show extreme differences in generated DSB/IRIF numbers, IRIF properties, and intensity of the background signal [35,36,59,60]. This can be attributed to the random character of damage induction, generation of DSBs that are repaired with unequal efficiencies, heterogeneous cell states, asynchronous repair of individual DSBs, and cells’ biological variability. Besides, IRIF parameters are influenced by the sample preparation too [30,61]. Confocal microscopy image data thus cannot be considered fully quantitative without a careful calibration and detailed knowledge on the exact experimental and biological behavior of the cell type studied. Simplistic intensity thresholding or approaches based on pre-defined parameters thus frequently fail, making the correct identification of IRIFs impossible. Accordingly, *The Second γH2AX-Assay Inter-Comparison Exercise* carried out in the framework of the European Biodosimetry Network (RENEB) [30], other available literature sources, and our experience have demonstrated that manual inspection of images by an expert eye still ensures more precise IRIF identification than automatic software algorithms. Nevertheless, visual quantification of IRIFs is extremely time demanding and difficult even for a trained eye. Unless all data are analyzed by a single observer, which is practically impossible, the results may suffer from dramatic variations [28]. Hence, the results obtained by different observers and/or labs can only be compared with extreme caution [30,62,63]. This unsatisfactory situation means that, without suitable software, the evaluation of large image datasets, as generated for instance in the case of mass radiation accidents, remains unrealistic. This problem strongly complicates also other practical (e.g., medical) applications and research.

Moreover, the information on architectural IRIF properties, such as the focus size, intensity, and shape, is left unexplored by the visual-only evaluation. The IRIF architecture has been recognized as an important factor regulating DSB repair processes and potentially participating in the decision-making for a particular repair mechanism (pathway) at individual DSB sites [36,37,55,64]. Architectural IRIF defects or their enhanced presence often appears in cells affected by cancer [35,36,65,66], precancerous syndromes [14,15,66] and, for instance, ageing [67]. Hence, improved ability of automatic software detection coupled with detailed characterization of IRIFs formed by individual repair proteins would be immediately recognized in numerous research fields as well as important practical areas of human activity related to DNA damage (e.g., medicine, radiation protection and space exploration).

Several strategies to segment IRIFs (usually γH2AX or 53BP1) have recently been published [68–72]. Focinator [68,73], FindFoci [69], the method proposed by Feng et al. [70], Foco [71], AutoFoci [72] or FocAn [74] represent the most important open-source attempts. Commercial software packages developed for microscopy image processing by microscopes providing companies are not focused on IRIFs specifically and the ongoing effort to develop new open-source IRIF analysis platforms clearly demonstrates that many important issues have not been solved satisfactorily.

To conclude, while automatic IRIF quantification (and nucleus segmentation) with only simple processing techniques, like thresholding, is inefficient due to variability of the fluorescence intensity and other IRIF parameters between the cells and experiments, huge amounts of image data related to emergency biodosimetric events or required to meet the research requirements preclude manual analysis of IRIFs. Even if there is sufficient manpower for this purpose, the results of different evaluators (even from the same lab) usually suffer from a strong subjective bias and are therefore hardly comparable. Moreover, IRIF parameters cannot be quantified except for the number.

In the present manuscript, we introduce DeepFoci, a novel robust software based on machine (deep)-learning strategies for fully automated identification and characterization of IRIFs formed by different repair proteins. The software overcomes serious shortcomings with IRIF detection described above and allows segmentation of cell nuclei and IRIFs with high fidelity, even in the case of challenging cell specimens of dramatically different quality as they appear in daily practice. The precision, specificity and reproducibility of the procedures are further enhanced by dual DSB labelling and colocalization analysis of two selected independent DSB markers [21,34]. At the same time, this allows studies on spatiotemporal interactions between IRIF proteins during the repair. The software has been successfully trained and tested on extremely challenging datasets based on tumor cell primary cultures and, in all cases, performed comparably to a careful, time-demanding manual analysis by an experienced expert.

## Methods

### Cells & Cell culturing

Following cells were used: (1) Normal and cancerous standard permanent cell lines represented by primary normal human dermal fibroblasts (NHDF, PromoCell, Heidelberg, Germany) isolated from the dermis of juvenile foreskin or adult skin and highly radioresistant U-87 glioblastoma cells (ATCC HTB-14, LGC Standards, United Kingdom), respectively. (2) Tumor and tumor-adjacent primary cell cultures isolated from patients with spinocellular head and neck tumors (histologically verified tumor and tumor-adjacent tissues). While the permanent cell lines represented biologically homogeneous and technically relatively easy samples, tumor cell primary cultures were involved as highly heterogeneous and challenging samples.

The protocol for patients’ primary culture isolation was described in [75]. The primary culture was cultivated in RPMI-1640 medium with the Pen/Strep antibiotic solution (PAA Laboratories GmbH, Austria) and 10 % fetal bovine serum, FBS (Biochrom, USA), at 37 °C and 5.0 % CO2 in a humidified atmosphere up to 50 % confluence. NHDF and U-87 were grown in Dulbecco’s modified essential medium (DMEM, Life Technologies) supplemented with 10% fetal calf serum (FCS) and a 1% gentamicin–glutamine solution (all reagents from Sigma-Aldrich).

### Irradiation

The cells were irradiated at the Institute of Biophysics of the Czech Academy of Sciences, Brno, Czech Republic in following schemes: (a) NHDF and U-87 cells were irradiated with increasing single γ-ray doses of 1, 2, or 4 Gy (D = 1 Gy/min) produced by Chisostat irradiator (^60^Co, Chirana, CR). (b) Patient-derived primary cultures were irradiated with a single dose of 2 Gy under the same irradiation conditions as the permanent cell lines. The cells were irradiated in the appropriate culturing medium at 37 °C, in a normal atmosphere. Irradiated cells were spatially (3D) fixed at indicated periods of time post-irradiation (PI), immuno-labelled, and visualized by confocal microscopy as described in the particular paragraphs below.

### Cell fixation and immunostaining

Aliquots of non-irradiated cells (0 min PI) and irradiated cell samples were washed in PBS and spatially (3D) fixed with 4% buffered paraformaldehyde for 10 min at RT at different periods of time post-irradiation (PI) —30 min, 8 h and 24 h PI. Subsequently, cells were permeabilized with 0.2% Triton X-100/PBS for 15 min and immune-labeled for IRIFs. Two combinations of primary antibodies were used for the immunofluorescence detection: anti-phospho-Histone H2AX (mouse, clone JBW301; Merck Millipore, Darmstadt, Germany, cat. no.: 05-636; 1:400) + anti-53BP1 (rabbit; Cell Signaling Technology, Danvers, MA, USA, cat. no.: 4937; 1:400), or anti-phospho-Histone H2AX (mouse; Merck Millipore; 1:400).

Among other ionizing radiation-induced (repair) foci (IRIFs) and DSB markers, γH2AX foci, 53BP1 foci were selected for the following purposes: γH2AX foci were used as a DSB marker in numerous studies and can point to changes of chromatin structure that appear at DSB sites during DSB repair. 53BP1 protein participates in DSB repair and the decision making process for non-homologous end joining (NHEJ) or homologous recombination (HR) at particular DSB sites. Since 53BP foci well colocalize with γH2AX foci and are formed with similar kinetics to these foci, 53BP1 and γH2AX foci are often co-detected to enhance the fidelity and reliability of DSB quantification [21,34].

The immunodetection procedure was described earlier [21,34]. Briefly, after the incubation with primary antibodies (overnight at 4°C), a mixture of secondary antibodies was applied for 1h (RT). Primary antibodies were visualized by the mixture of FITC-conjugated donkey anti-mouse and Cy3-conjugated donkey anti-rabbit (both Jackson Immuno Research Laboratories, West Grove, PA, cat. no.: 715-095-150 and 711-165-152) applied in 1:100 and 1:200 dilutions, respectively (30 min incubation at RT in dark). Alternatively, anti-mouse Alexa Fluor 647 and anti-rabbit Alexa Fluor 568 (ThermoFisher Scientific) secondary antibodies were used, which are directly compatible with both confocal microscopy and SMLM. The antibodies were diluted in sterile Donkey serum (1:400 and 1:200, respectively; cat. No.: P30-0101, Pan Biotech GmBH) and applied to the cells for 30 min (RT, in the dark). After incubation, the cells were washed 3x in 1× PBS for 5 min.

The cell nuclei were stained with DAPI (5 min at RT) provided as Duolink *In Situ* Mounting Medium with DAPI (DUO82040; Sigma-Aldrich; now Merck, Darmstadt, Germany) and diluted to the concentration of 1:20.000 [76]. Afterwards, the slides with cells were washed 3x in 1× PBS for 5 min each. Finally, the coverslips were air-dried, and the cells were embedded in ProLong Gold (ThermoFisher Scientific). The Prolong Gold was left to polymerize for 24 hours in the dark at RT. After complete polymerization, the slides were sealed with nail polish and stored in the dark at 4° C. Alternatively, nuclear chromatin was counterstained with 1 μM TO-PRO-3 (Molecular Probes, Eugene, OR) in 2× saline sodium citrate (SSC), prepared fresh from a stock solution. After brief washing in 2× SSC, Vectashield medium (Vector Laboratories, Burlington, Canada) was used for the final mounting of the samples.

### Confocal microscopy

Leica DM RXA microscope equipped with DMSTC motorized stage, piezo z-movement, MicroMax CCD camera, CSU-10 confocal unit and 488, 562, and 714 nm laser diodes with AOTF was used for acquiring detailed cell images with 100x oil immersion Plan Fluotar lens, NA 1.3) with a Z step size of 0.3 μm. The equipment was controlled by the Acquiarium software developed by [77]. The resulting images are 90.0×67.2×15 μm xyz (1392×1040×50 px). Usually, forty serial optical sections were captured at 0.25 μm intervals along the z-axis. The R-G-B exposure times and the room/device temperature were optimized for individual samples to provide optimal images.

### Datasets for software analyses

The training/validation/testing datasets was based on patient-derived primary cell cultures prepared from spinocellular tumors and morphologically normal tissues adjacent to the tumor taken from patients suffering from head and neck cancer. The dataset was divided into two subsets: one for training, validation and testing the nucleus segmentation (237/10/30 fields of view (FOVs), respectively) and one for training, validation and testing the focus segmentation (239/60/100 FOVs). The dataset consisted of several cell types: a) tumor cells, b) tumor-associated fibroblasts, and c) cells from morphologically normal tissues. All cell types were fixed at different periods of time (0 (non-irradiated control), 0.5, 8 or 24 h PI) after exposure to 2 Gy of γ-rays. The representation of cells in two subsets with respect to the cell type and post-irradiation time (i.e., DSB repair duration) was random.

The evaluation dataset was used to assess the robustness of segmentation procedures. It was composed of multiple types of differently treated cells in order to represent a highly challenging dataset maximally reflecting high biological and technical variability between samples, as it may appear in research or clinical practice. The dataset contained a) mesenchymal NHDF fibroblasts coming from a standard permanent cell line, b) radioresistant U-87 glioblastoma cells coming from a standard permanent cell line, c) tumor cells (CD90^-^) and tumor-associated fibroblasts (CD90^+^) prepared as a primary culture from a spinocellular tumors of patients (different from dataset 1) suffering from a head and neck cancer, and d) cells prepared as primary cultures from morphologically normal tissue adjacent to tumors of involved head and neck cancer patients. NHDF and U-87 cells received 0, 1, 2 or 4 Gy of γ-rays and were fixed at 0.5 PI, while the primary cultures were only exposed to the dose of 2 Gy (for a limited amount of the cell material) and fixed at 0 (non-irradiated control), 0.5, 8 or 24 h post-irradiation times.

### Ground truth generation

Manual annotation of nuclei and IRIFs is required for CNN training and performance evaluation. As both nucleus segmentation and IRIF detection are performed in 3D, manual labelling is problematic with available labelling tools. For this reason, a customized labelling GUI was created in Matlab, enabling easier 3D data labelling. The tool for the nucleus segmentation is based on the pre-segmentation of an image with SLIC superpixels approach [78], where superpixels are then labelled in GUI by the user. For the detection training, IRIFs were pre-detected with the same algorithm as for the final segmentation, where CNN is replaced by a simple local maxima detector applied on the colocalization image. This detector is set to high sensitivity to capture all potential IRIFs. The user then selects real foci from IRIF proposals in the 2D projection image, where its 3D coordinates are taken from the detector. Nuclei and IRIFs for training were manually annotated in 3D by one expert. IRIFs for testing of the algorithm were labelled in 2D projection.

### Evaluation metrics

To evaluate the accuracy of nucleus segmentation in 3D, SEG score (object-wise Intersection over Union (IoU)) was used [79]. To calculate SEG, IoU (also known as Jaccard index) is needed for the calculation. IoU is defined as equation (1)

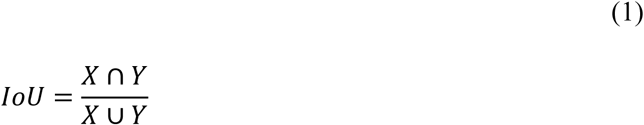

where X and Y are manually annotated segmentation mask and predicted segmentation mask, respectively. For every manually annotated object, the segmented object with the largest IoU is found. Next, average IoU of all manually annotated objects is calculated. If IoU for any manually annotated object is smaller than 0.5, then IoU for this object is set to 0. This ensures that each manually annotated object can be paired with only one segmented object. The resulting SEG is an average of IoUs for all manually annotated objects. To evaluate the detection accuracy of individual IRIFs, Dice coefficient (F1-score) was used. Dice coefficient is defined as equation (2)

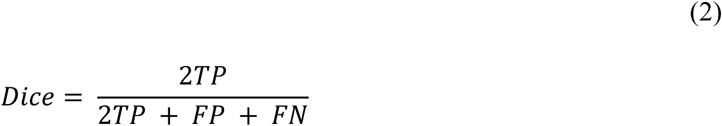

where TP is the number of true positive IRIFs, FP is the number of false positives and FN is the number of false negatives.

### Ethical declarations

The study was conducted in accord with the Helsinki Declaration of 1964 and all subsequent revisions thereof. It was approved by the ethical committee of St. Anne’s Faculty Hospital, Brno.

## Results

To overcome important shortcomings of currently available procedures for the DNA repair focus (IRIF) analysis, we have developed DeepFoci, a novel software tool based on artificial neural networks and deep-learning that allows fully automated detection, quantification and analysis of these structures in the context of their natural environment, i.e., within the architecture of the cell nucleus (chromatin). DeepFoci is written in Matlab and is primarily focused on precise 3D segmentation of IRIFs and cell nuclei in large datasets of confocal microscopy images and consequent analysis of recorded data. To fully benefit from the software abilities, IRIF visualization with fluorescently-tagged antibodies against two different DSB markers is applied. This dual labelling improves the precision of the DSB quantification and, at the same time, allows to study the spatiotemporal relationship between γH2AX (or other epigenetic modifications), repair proteins of interest, and chromatin architecture/function at individual DSB sites. Nevertheless, the analysis based on a single IRIF marker (e.g., γH2AX) staining is also possible if preferred by the character of an experiment or practical situation. In the present manuscript, γH2AX and 53BP1 were selected as the IRIF markers—γH2AX because of its widespread usage for this purpose and 53BP1 protein for its involvement in both main DSB repair pathways (the non-homologous end-joining (NHEJ) and homologous recombination (HR)) [80,81]. Moreover, γH2AX and 53BP1 foci differ in their morphological features but share a similar formation-decomposition kinetics and extensively colocalize with each other.

Images were preprocessed with a 5×5×1 median filter, Gaussian filter with sigma 1 and image normalization, where values between 0.0001 and 99.999 percentile were transformed to the range of 0– 1. The proposed IRIF focus analysis method consists of three main steps (see Fig. 1): (1) initial instance segmentation of single nuclei with Convolutional Neural Network (CNN), (2) detection of individual IRIF foci again with CNN and (3) segmentation of detected foci with Maximally Stable Extremal Region (MSER) algorithm.

**Figure 1.**
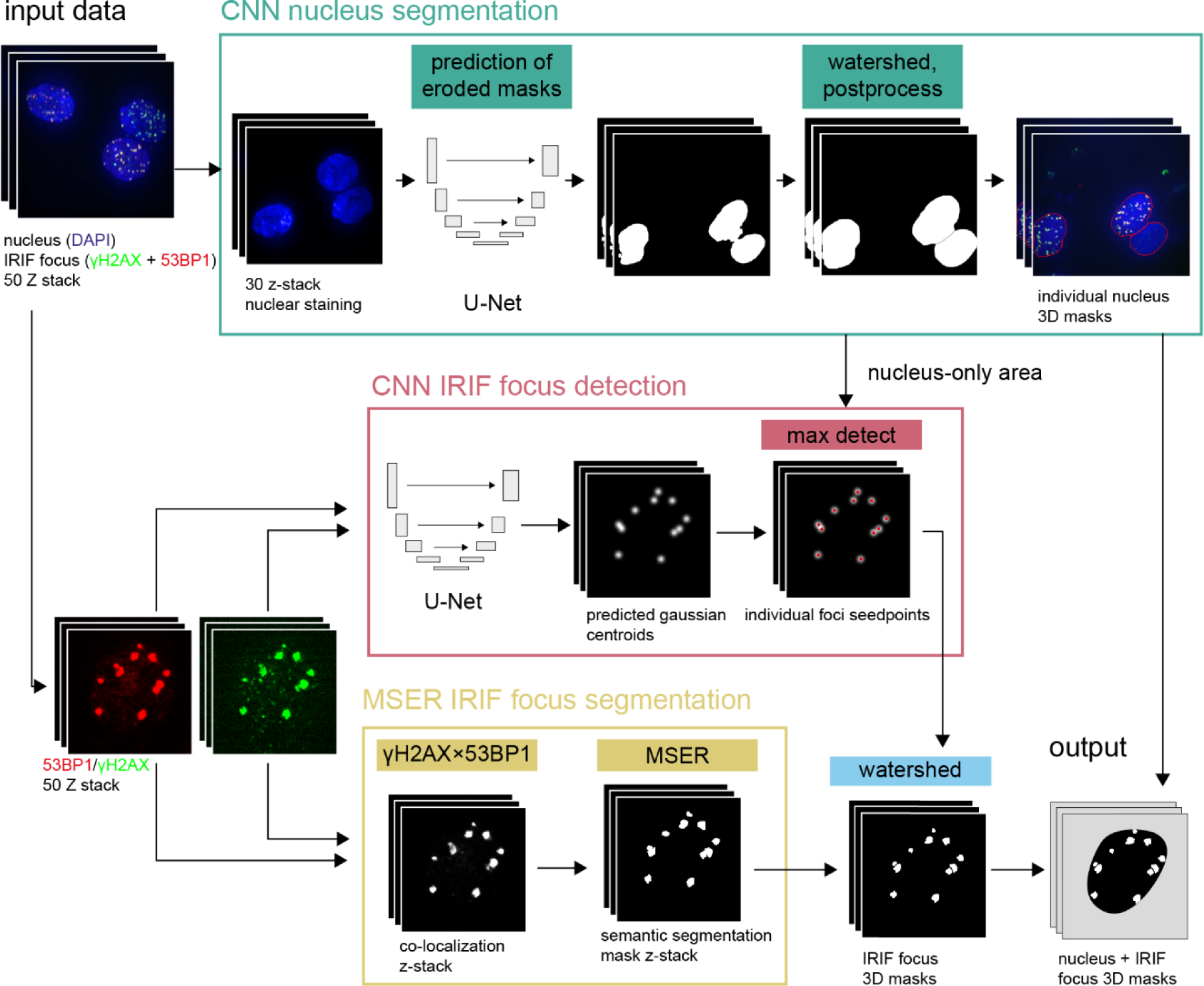
Block diagram of IRIF detection. 3-channel images are used for a network input: one for nuclear staining and two for IRIF staining, as exemplified by DAPI staining of the nuclei (blue) and immunodetection of γH2AX (green) and 53BP1 (red) IRIFs. The process is divided into three steps: First, 3D nucleus masks are created using U-Net CNN from the channel for the nucleus staining. Second, individual IRIF foci are detected with a convolutional neuronal network (CNN). Third, individual foci are segmented from a multiplied z-stack composed of the two channels for IRIFs utilizing maximally stable extremal region detector (MSER). The output of these three steps is finally merged into nucleus/IRIF 3D masks.

### Nucleus segmentation

Image-to-image encoder-decoder CNN with the U-Net topology [82] had proved to be very powerful for biomedical image segmentation. However, it produces just foreground-background (semantic) segmentation in most standard cases. The standard U-Net architecture will not ensure single nucleus separation as every error in the boundary pixels would result in the connection of neighboring nuclei into one segmented object. Successful separation of individual nuclei was achieved by modifying the network in order to predict the eroded binary masks. However, this can still lead to incomplete separation of individual nuclei due to prediction errors on the boundary between nuclei. For this reason, 3D CNN prediction is followed by the distance transform (DT) and watershed segmentation (applied to the negative of the DT image) to separate the touching nuclei [83]. Moreover, the distance transform image is processed with the *h*-maxima transform and grayscale dilatation, which prevents over-segmentation. This step removes the maxima that are close to each other and separated by an insufficient decrease in the image intensity. The minimal distance between the maxima is controlled by the radius of the structuring element and the minimal image intensity decrease defined by the *h* parameter of the *h*-maxima transform. Afterwards, the resulting image is dilated to compensate for the initial erosion of ground truth masks. In order to prevent nucleus merging, the dilatation is performed sequentially for individual nuclei. Both *h* parameter and minimal distance between the maxima were optimized with grid search on a validation set.

### IRIF detection

Similarly, 3D U-Net was applied for the detection of individual IRIF foci. In this case, an image with the 3D Gaussian function overlaid over the position of each IRIF centroid represented the ground truth for CNN training. Thus, the 3D CNN predict an estimation of possible foci in the form of Gaussian function, which is further post-processed. Individual foci were detected using the maxima detect—the local maxima with a value above the threshold were considered as detected foci and the *h*-maxima transform and grayscale dilatation were utilized to prevent multiple detections of the same IRIFs due to inaccurate prediction of CNN. Both the threshold and the *h* parameter were optimized with the grid search on a validation set.

### IRIF segmentation

The Maximally Stable Extremal Regions (MSER) [84] is a segmentation technique that is generally very robust to illumination changes and therefore suitable for the segmentation of fluorescence microscopy images of varying intensity. Extremal regions of an image are defined as the connected components of a thresholded image. MSER produced stable extremal regions of the image, which are stable in the sense of the volume variation w.r.t. changes of the threshold. The minimal allowed stability of the extracted region can be set with two parameters—the threshold step and the maximal relative volume change with this step.

This IRIF segmentation approach was applied to the colocalization image created as a multiplication of the two IRIF channels (as represented by γH2AX and 53BP1 signals in here). Using MSER, multiple segmentation variants of increasing size for every IRIF can be generated. Its size was restricted to the maximal IRIF volume. Of the segmentation variants, the largest one is selected as a final segmentation mask. The IRIF segmentation produced by MSER was then combined with the U-Net IRIF detection employing the seeded watershed transform. The seeded watershed transform [85] was then applied on the colocalization images, where the outputs of the IRIF detection served as the seeds and MSER served as foreground mask.

### Implementation details

Matlab R2019b with Image processing and Deep Learning Toolboxes and VLFeat library (for 3D MSER) [86] was used. The 3D U-Net network [82] with 16 filters in the first layer was employed for both the nucleus segmentation and IRIF detection setting. For the nucleus and IRIF detection, only the loss functions were different—the Dice loss [87] for nucleus segmentation and the Mean-Squared error loss for IRIF detection. For higher computational efficiency and GPU memory limit of the nucleus segmentation and IRIF detection, the image volumes were downscaled in X-Y dimensions by a factor of two (505×681×50 px). Augmentation with a selection of random patches (96×96×50), random flips, and multiplication of image pixels by a random value between 0.6 and 1.4 was used.

### Nucleus segmentation evaluation

The SEG instance segmentation measure [79] was adapted for the segmentation of nuclei. Two nuclei were considered matching if the IoU was equal or greater than 0.5. Each ground truth nucleus was included exactly once to prevent assignment to multiple nuclei. The nucleus segmentation was tested on a dataset consisting of 30 FOVs annotated by a single expert, with the same tool as used for the generation of the training data. The proposed method achieved a median SEG score of 0.82 (median over FOVs). The representative FOV with the SEG of 0.80 and the distribution of SEG values (a histogram for all FOVs) is shown in Fig. 2a and 2b, respectively.

**Figure 2.**
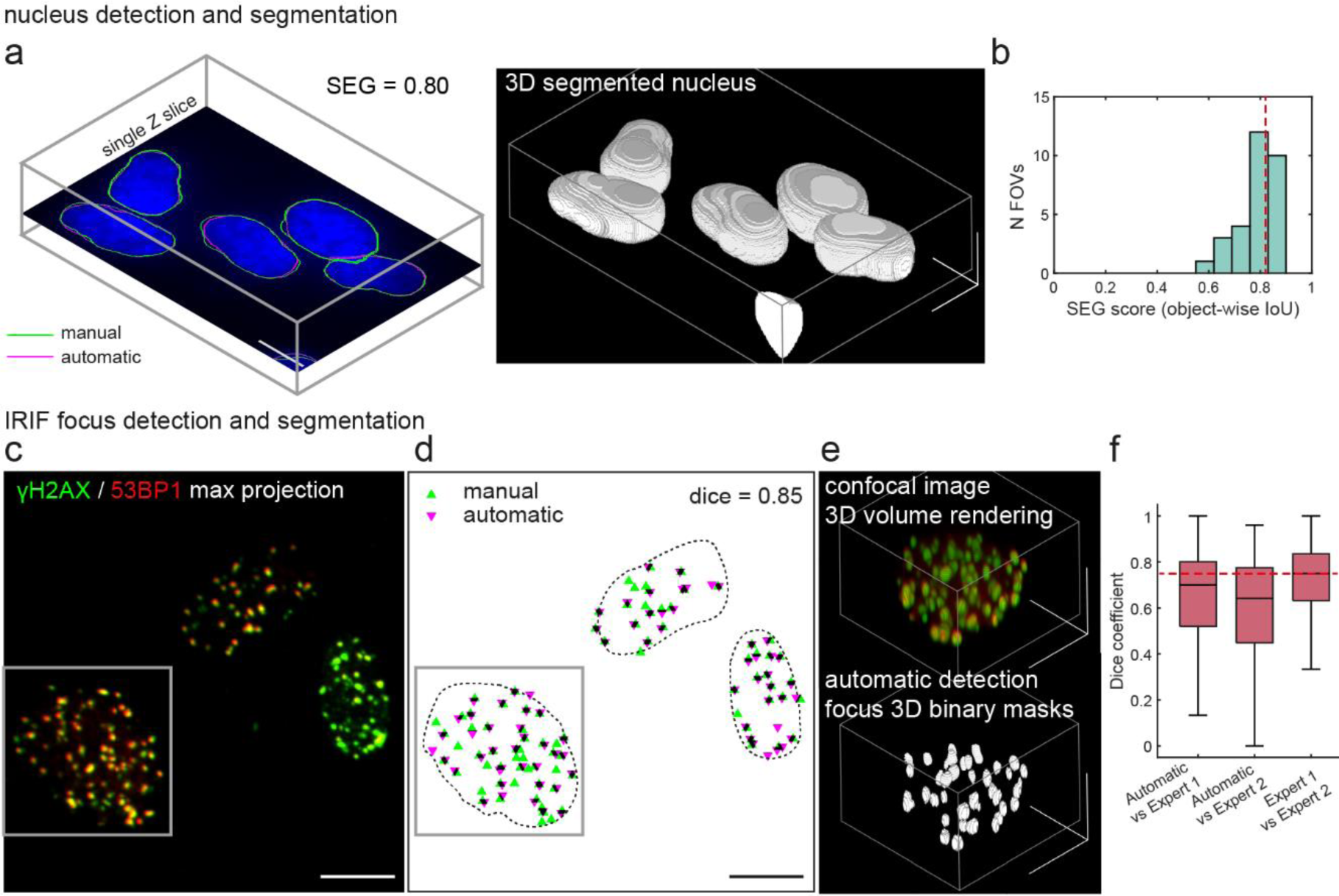
The results of nucleus segmentation and IRIF detection. **a-b** The nucleus segmentation performance; **a**. Comparison of the automatic and manual nucleus segmentation. Segmentation results for a single field of view (FOV), single Z slice, NHDF cells, and DAPI staining is shown (left), together with 3D reconstruction (right). **b**. Histogram of the SEG score for the nucleus segmentation for 30 segmented FOVs used for testing. Red line indicates median SEG of all FOVs. **c-f** The IRIF focus segmentation performance. **c**. IRIFs after 2 Gy γ-ray exposure, 8 h post-irradiation, oropharyngeal squamous cancer cells, γH2AX/53BP1-staining, max projection, 100x magnification. **d** Comparison of manual annotation and DeepFoci detection result. **e**. Top - 3D confocal data of the detail indicated by a grey square in 2c; bottom - binary masks detected by proposed CNN. **f**. The IRIF focus detection performance, comparison of the automatic result with two manual annotations by experts shown as median, IQR and min-max in 1.5 IQR. Red line indicates median Dice coefficient of IRIF detection between two experts. Scale bar in all FOVs indicates 10 μm.

### IRIF detection evaluation

The accuracy of automated IRIF detection was compared to manual annotation performed by two experienced experts on the maximum-projection images. The Dice coefficient served as the IRIF detection accuracy metric (see Methods). The IRIFs detected by the proposed software procedures and annotated manually, respectively, were considered as mutually matching if their centroids were closer than 1.95 μm (30 px), a value corresponding to maximal dispersion of manual annotations between experts. The IRIF detection performance using DeepFoci is presented in Fig 2c-e. Manual annotation by two experts enabled not only to evaluate the IRIF detection by DeepFoci itself, but also estimate the minimum difference that can be experienced between experts. The difference between manual expert annotations provided a median Dice coefficient equal to 0.75. The discrepancies between the automated IRIF detection/segmentation by DeepFoci and manual annotations by either of the experts were close to the variability between experts, with Dice coefficients 0.64 for the expert 1 and 0.70 for the expert 2 (Fig. 2f). An example of the automated IRIF detection and segmentation is presented in Figs. 2c-e.

### Method verification and practical applications

The DeepFoci performance was compared with two recently published tools for IRIF counting: FocAn [74] and AutoFoci [72] and with CellProfiler – a universal particle analysis software. For AutoFoci and FocAn only IRIF segmentation part (without nucleus segmentation) was applied. For 2D methods (AutoFoci and CellProfiler), the maximum intensity projection images were used. CellProfiler and FocAn are designed for single IRIF fluorescence staining, thus, the procedures were applied to the colocalization images (obtained by multiplication of the two IRIF channels), because this variant achieved the best results. Parameters of these methods were optimized using the grid-search. For AutoFoci, the most reliable values were searched and set up for *the object evaluation parameter threshold, minimal focus distance* and *top-hat structuring element radius*. For FocAn, *the minimal focus distance, threshold value* and *neighborhood size of adaptive threshold* were optimized. CellProfiler employed a simple pipeline (based on [69]) with *EnhanceOrSuppressFeatures–Enhance Speckles* option and with *IdentifyPrimaryObjects–Otsu method* and distance local maxima suppression. *The minimal focus distance* and the parameter of *Enhance Speckles* were tuned.

Based on the performance on our challenging testing dataset, composed of heterogeneous primary cell cultures derived from squamous head and neck cancer patients’ tumors (see Methods for details), the highest accuracy among all compared software detection methods was achieved with DeepFoci. The median Dice coefficients for FocAn, AutoFoci, CellProfiler and DeepFoci were 0.22, 0.38, 0.49 and 0.67, respectively (Fig.3a).

To further compare DeepFoci with these already-available software approaches, the correlation between the manually annotated and automatically detected IRIFs was evaluated for nuclei with various IRIF counts, ranging from 0 or only few in non-irradiated controls to several dozens in cells fixed at 1 h post-irradiation. To cover all stages of DSB repair that dramatically differ in the number of IRIFs per nucleus and IRIF parameters, primary tumor cultures were fixed at different periods of time till 24 h post-irradiation (PI). Specifically, the fixation times were selected to test the software ability to quantify a) large amounts of morphologically variable IRIFs at the time of their maximum appearance in nuclei (0.5 h PI), b) middle amounts of large but differently diffused late IRIFs (8 h PI) and c) low only amount of few persistent IRIFs (“irreparable” DSBs) in cells that almost accomplished repair (24 h PI). Non-irradiated cells (0 min PI) served as the negative controls with none or only few naturally occurring IRIFs. Among all software approaches (Fig 3b), the IRIF numbers detected by DeepFoci best correlated (had the most linear dependence) with the values obtained for the same corresponding nuclei by manual expert analysis (average values for individual experts are plotted at Fig. 3b). Moreover, unlike other tested methods, DeepFoci retained its sensitivity and fidelity at the same time over the whole scale of possible IRIF amounts.

**Figure 3.**
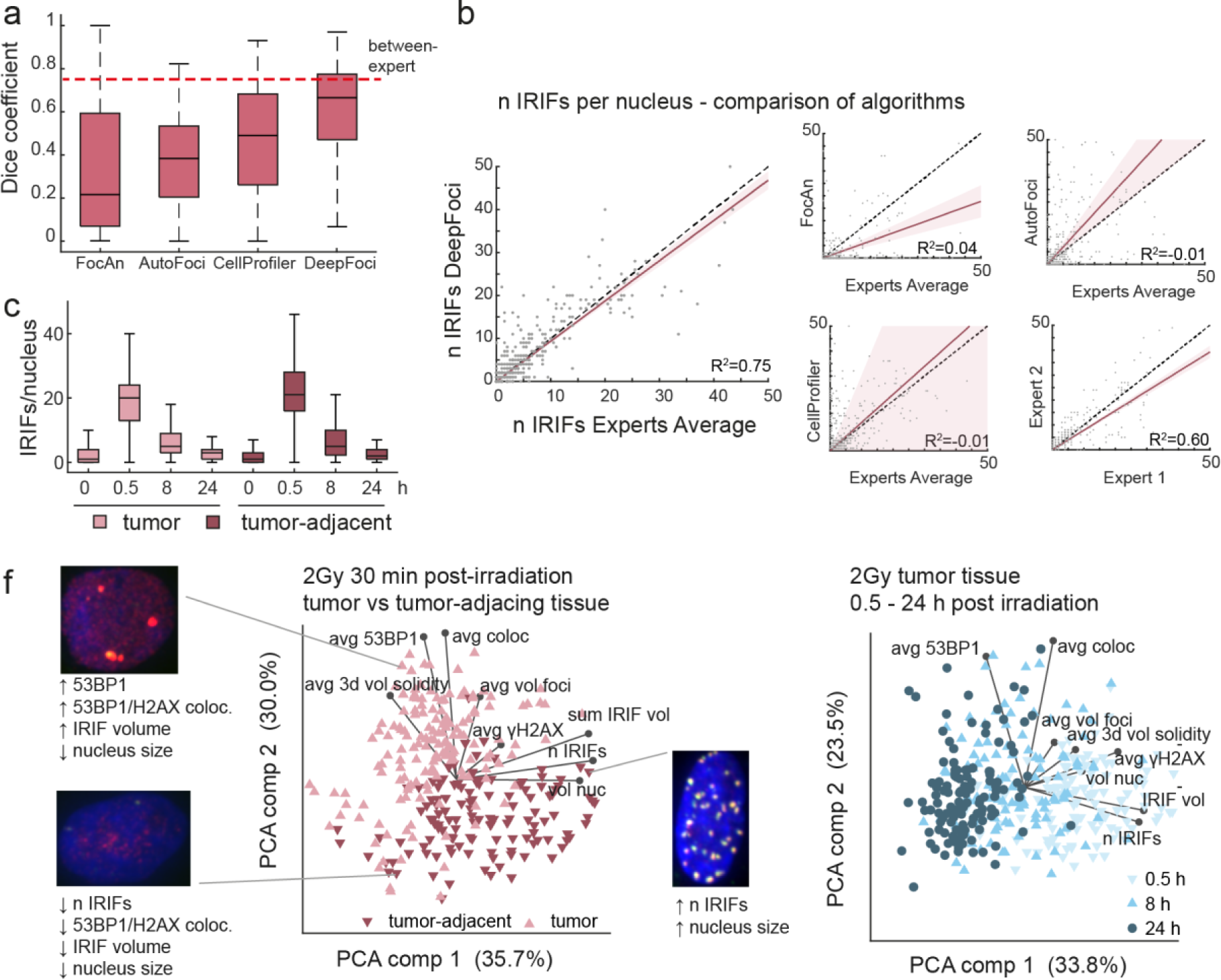
DeepFoci performance. **a**. Performance comparison for published IRIF-detecting approaches and DeepFoci. Results obtained for the challenging dataset based on the head and neck squamous cell cancer primary cultures are shown. Red dashed line indicates the conformity (Dice coefficient) between experts. **b**. The correlation between automatically and manually detected (averaged results for two expert annotations are plotted) IRIF numbers compared for DeepFoci and all other tested software methods. **c**. The DSB repair kinetics determined on the basis of the average IRIF numbers per nucleus detected by DeepFoci at different periods of time (0 min – 24 h) post-irradiation. The repair behavior is compared for normal fibroblasts (NHDF) and highly radioresistant U-87 tumor cells. **d**. The principal component analysis biplot showing the separation of the tumor-adjacent tissue cells and tumor tissues cells (left) and the separation of tumor cells fixed at different post-irradiation times (right). The inserts show representative nuclei with IRIFs for revealed cell subgroups (categories).

To demonstrate practical usability of DeepFoci and further test its performance, DSB repair kinetics and IRIF morphology was compared for the head and neck squamous cell cancer primary cultures and tumor-adjacent cultures, using additional FOVs that were not involved in the training procedure. For all the tumor cell primary cultures and tumor-adjacent non-tumorous primary cultures, the IRIF numbers peaked at 0.5 h PI, which was followed by a significant drop at 8 h PI and persistence of only few IRIFs/nucleus at 24 h PI (Fig. 3c). Such a repair profile (repair kinetics) corresponds well with the profile that can be expected for cells exposed to 2 Gy gamma radiation [13], i.e., the conditions used in the present work.

Besides the number of IRIFs per nucleus, analyzed as the only parameter in most studies, a wide spectrum of additional parameters can be extracted by DeepFoci, including the focus intensity in two color channels, intensity of chromatin staining at the IRIF site and extent/character of all color channel colocalization. Furthermore, 3D morphometric features of IRIFs and nuclei—such as their volume, solidity, and circularity—were possible to measure. The principal component analysis biplot (Fig 3c) shows the interdependence between these parameters and examples of multi-parameter classification/categorization of IRIFs. The graphs demonstrate that the separation of cell groups of interest—as plotted for tumor vs. tumor-adjacent (normal) tissue cells (left) or cells left to repair DSBs for various PI times (right)—is much better compared to separation solely based on the IRIF numbers. This result demonstrates that the IRIF parameters could be mutually interdependent in a complex way, so that their joint consideration may allow categorization of cell groups even in cases when they cannot be separated solely on the basis of IRIF numbers. In our head and neck squamous cell cancer dataset, the tumor tissue cells and tumor-adjacent tissue cells with similar average IRIF counts per nucleus were distinguished when the morphology of IRIFs (e.g., the average 3D solidity) and intensity of 53BP1 foci were taken into account. DeepFoci analysis also revealed several distinct cell groups within the same dataset that corresponded to cells fixed at different periods of time post-irradiation (Fig 3c, right). In this case, especially the inclusion of γH2AX focus intensity emerged as an important parameter, in addition to the extent of γH2AX and 53BP1 mutual colocalization. This is in accordance with the known fact that 53BP1 binds to γH2AX early after DSB induction and dissociates when the damage is repaired, which is also accompanied by γH2AX dephosphorylation. The degree of colocalization between γH2AX and repair proteins thus proved useful for separation of cells in different phases of the repair process and, in some cases, also separation of normal and tumor cells.

Finally, DeepFoci algorithm was verified on a dataset composed of permanent cell lines of normal human skin fibroblasts (NHDF) and U-87 glioblastoma cells. U-87 cells are derived from a radioresistant brain tumor that is treated by radiotherapy while NHDF fibroblasts are normal (non-transformed) cells with relatively lower radioresistance that are always exposed to radiation during radiotherapy or in the event of an radiation accident. For these differences between NHDF and U-87 cells and their different origin, cell-type specific IRIF morphology and repair dynamics can be expected (as already reported in [36]). The cells of both types were exposed to γ-ray doses ranging from 1 to 4 Gy and fixed at different periods of time post-irradiation. Figure 4 shows the results for 30 min post-irradiation, i.e., the post-irradiation time when a mixture of well-developed and immature IRIFs can be seen. The analysis by DeepFoci closely approached the precision of manual analysis under all conditions tested (times PI, radiation doses, and cell types) and the maximum average numbers of IRIFs per nucleus detected by DeepFoci felt well within the interval of values (about 14 to 25) reported in most studies for given conditions and various cell types [14–16,18,20,28,34,40,41,76,88,89].

**Figure 4.**
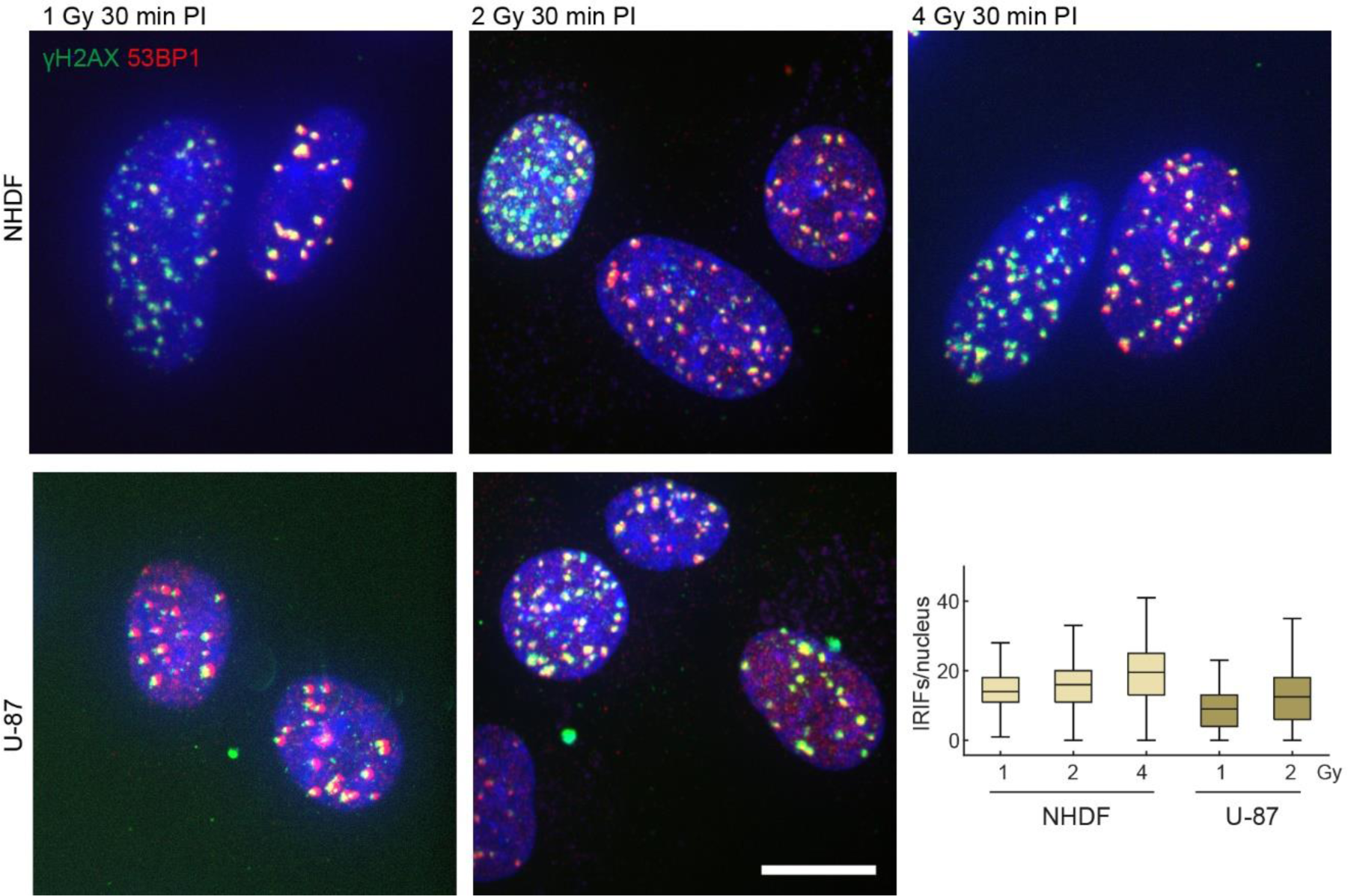
DeepFoci-based detection of IRIFs compared for different radiation doses and cell lines with varying sensitivity to γ-rays. IRIF numbers detected by DeepFoci in normal fibroblasts (NHDF) and radiotherapy-resistant glioblastoma cell line, U-87, after irradiation with increasing γ-ray doses and fixation at 30 min post-irradiation (PI). Representative FOVs of NHDF and U-87 are displayed. Scale bar indicates 10 μm.

## Discussion

The discovery of IRIFs has been a milestone in radiobiology and medical research. Currently, monitoring of IRIFs represents the most sensitive and versatile method to quantify DSBs and study repair protein interactions and epigenetic modifications at individual DSB sites, within the natural environment of the cell nucleus and in time. Immunofluorescence microscopy provides the most complex information on IRIFs so it has found irreplaceable applications in biodosimetry and research. However, robust, precise and reproducible identification of IRIFs still represents unsatisfactorily solved task, even in the current era of advanced image analysis technologies. *The Second γH2AX-Assay Inter-Comparison Exercise* carried out in the framework of the European Biodosimetry Network (RENEB) [30] and our experience have shown that a visual (manual) identification of IRIFs still far overpowers the performance of any software package in terms of precision. On the other hand, the manual quantification of IRIFs by a single evaluator is usually unrealistic for extreme time demands. Cooperation of more evaluators does not help, unless they are intensively (i.e., for a long time) trained to reduce inter-expert biases in IRIF identification (see Fig. 2f). Equally frustrating is the principal inability of manual analysis to extract IRIF parameters that are of fundamental interest in DNA damage and repair research and may be of practical relevance too. This precarious situation strongly limits analyses of larger datasets, results comparison both within and between labs and future progress.

Automated (software) IRIF segmentation is mostly challenged by a tremendous variability of these structures in all their parameters. Mainly the amount of fluorescence can vary a lot between samples. Simple thresholding-based strategies thus provide acceptably accurate and reproducible results only in specific situations, e.g., when the same cell type (e.g. normal lymphocytes) is repeatedly analyzed using a well-optimized staining procedure. Most available methods therefore use some adaptive thresholding or image standardization; however, this leads to the dependence of the detection sensitivity on the number of IRIFs in the nucleus (Figure 2b). In principle, it remains impossible to set up a threshold parameter universally so that all IRIFs (early, mature, and late) can be correctly recognized and segmented. Specific settings to detect IRIF parameters are often necessary also for individual datasets. Particularly problematic are samples with little or no IRIFs (non-irradiated controls or cells that already accomplished repair, etc.), in which large numbers of false positive IRIFs are usually detected as a result of automatic thresholding or image standardization. Of the reports on IRIF-detecting algorithms, the ones on Foco [71] or Focinator [68,73] (introduced below) do not disclose the results for control samples. In the paper on FocAn, the controls (time point 0) are included but show unrealistically high IRIF numbers [74].

Several strategies trying to segment IRIFs (mostly γH2AX or 53BP1 foci) have recently been published [68–72]. Focinator [68,73] is a simple ImageJ macro enabling thresholding, maxima detection, and filtering based on the size and circularity. FindFoci [69] represents an ImageJ plugin that detects IRIFs as the local maxima. Focus regions are segmented with the downhill gradient algorithm and the proposed foci are eventually filtered out with specified parameters, which can be trained on a few labelled images. However, the procedure is suitable just for single-channel (γH2AX) labelling so that spatiotemporal interactions between repair proteins or repair proteins and chromatin within the IRIF or the cell nucleus cannot be studied. Feng et al. [70] use rather simplistic fuzzy c-means clustering for IRIF detection, which produces noisy and mutually incomparable results if IRIF amounts in nuclei vary to a higher degree. This shortcoming thus seriously complicates even basic analyses of DSB repair kinetics (IRIF number changes in post-damage time) as IRIF numbers may be very high after DNA damage induction (e.g., irradiation) but decrease to zero with repair time. Foco [71] presents an interesting pipeline for the nucleus and IRIF segmentation; however, the nucleus segmentation is based on the intensity thresholding, which we demonstrated to be insufficiently robust for our datasets. AutoFoci [72]—an advanced high-throughput algorithm—extracts several features from each IRIF and finds the most reliable one to distinguish between the IRIFs and noise. FocAn [74] is the only available 3D IRIF detection method implemented as an ImageJ macro; however, it is based on simple adaptive thresholding followed by maxima detection. CellProfiler offers a universal particle detection algorithm, where customized IRIF detection pipelines can be developed (pipeline from [69] was tested in this paper).

According to our experience [13], the difficulty with simple thresholding methods can be especially strongly experienced in patient-derived primary cultures that are characteristic by their high heterogeneity. Cells obtained from different patients show, by nature, dramatic differences in IRIF parameters and may even unpredictably react to the same staining protocol (the staining procedure optimization for particular patient samples is usually not possible due to a material lack and/or time demands). This was the reason why we included tumor cell primary cultures from different patients in our training and testing datasets.

The main motivation for this work was to explore whether the obstacles with IRIF detection and segmentation in confocal datasets could be overcome by employing deep learning. We aimed at enabling an unbiased analysis of large datasets in a timely manner, thereby allowing the realization of complex research studies, effective medical triage (biodosimetry) in the events of mass-casualty radiological incidents, and result comparison between samples and laboratories. In the present manuscript, we have introduced DeepFoci, a novel robust software based on deep learning for fully automated identification and morphometric characterization of IRIFs formed by different repair proteins in 3D. The software has been designed to overcome current limitations of fluorescence image analysis and allow segmentation of cell nuclei and IRIFs with high fidelity, even in the case of challenging cell specimens of dramatically different quality as they appear in daily practice. The results confirmed our idea that the precision, specificity and reproducibility of the procedures can be significantly enhanced by dual DSB labelling and colocalization analysis of two selected independent DSB markers [21,34]. At the same time, this strategy allowed us to analyze the recruitment of repair proteins into IRIFs and follow their spatiotemporal interactions during the repair. These achievements are crucial for both practical (e.g. clinical) and research applications.

It has been well documented (see also Fig. 3e) that individual IRIFs differ quite dramatically in their shapes, sizes, border sharpness and intensities. This variability appears between a) cell types [36], b) cell cultures (especially tumor cell primary cultures) and c) individual cells. Besides, it is also problematic to determine and define simple parameters that will optimally separate individual IRIFs within IRIF clusters. Using DeepFoci, multiple IRIF parameters were computed, which allowed multi-parametric IRIF categorization and thus more accurate recognition of different cell/patient groups and/or repair stages (Fig. 3f). The results indicated that morphometric parameters of IRIFs (such as the 3D solidity) as well as the extent and character of γH2AX and 53BP1 colocalization do change between cell types and post-irradiation periods. In turn, we show that this information, extracted by DeepFoci, can be used to further refine identification and categorization of different cell classes or (pre)malignant subclones. For instance, as compared with normal human skin fibroblasts, a lower degree of 53BP1 colocalization with γH2AX has been discovered at the nanoscale in U-87 glioblastoma cells [36,64], which are highly radioresistant. Here we show that cell-type differences in γH2AX and the repair protein colocalization can also be observed at the microscale and may point to important differences in DSB induction and repair between different (normal vs. tumor, radiosensitive vs. radioresistant, etc.) cells. The extent of colocalization also depends on the functional IRIF status. For instance, γH2AX marking unrepairable, still unrejoined DSBs could be expected to intensively colocalize with 53BP1 repair protein. On the other hand, IRIFs at sites of DSBs that were already ligated but where the reconstitution of the original chromatin architecture failed (epimutations [90]) may be solely decorated by γH2AX. Non-colocalizing 53BP1 foci can be induced by replication stress and were proposed to protect chromosomal fragile sites and DSBs that are transferred unrepaired to the next cell cycle [91]. Hence, DeepFoci broadens applicability of IRIFs as clinical biomarkers and makes possible to study DSB repair mechanisms (and their defects) at individual DSB sites.

DeepFoci was able to identify and quantify the number of IRIF foci with higher accuracy compared to CellProfiler [92], AutoFoci [72] or FocAn [74] and was of comparable precision to a careful manual analysis performed by a single experienced expert (Fig. 2f, 3a). The problems with the IRIF detection outlined in the previous paragraphs were circumvented by the introduction of a robust 3D segmentation technique based on the standard U-Net for binary segmentation. The main modifications involve the application of erosion on segmentation masks and subsequent post-processing of binary predictions, which lead to correct separation of individual nuclei. Similarly, the prediction of individual IRIFs via 3D Gaussian functions with specific postprocessing provided precise IRIF detection, which was close to human accuracy. As well, MSER showed to be a fast and powerful method for the 3D segmentation of IRIFs, producing very precise segmentation results with robust tolerance to different IRIF intensities. With these improvements, the method proved to achieve satisfactory segmentation of both nuclei and IRIFs. The main advantage of our method is, that it is robust against changes in the image intensity. It also uses the same U-Net architecture for both the nucleus segmentation and IRIF detection, which reduces its implementation complexity. In contrast to manual IRIF counting, the developed method is fast and automatic, and it provides the possibility to extract many other IRIF features besides the IRIF count, e.g., the mean intensity, size, and solidity. Compared to available automation attempts, DeepFoci is trainable and utilizes advanced deep-learning algorithms. This fundamental advantage lead to much better results than the methods based on the thresholding and maxima detection approaches. Moreover, the proposed approach operates on 3D samples. The only other available 3D method is FocAn [74]; however, it utilizes very simple IRIF detection approaches and provided unsatisfactory results on our challenging datasets.

Importantly, due to the learning-based nature of the implemented methods, the proposed algorithm offers an extensive room for application modifications and can be easily adapted for specific requirements of different laboratories, where it can be simply re-trained for a different type of data. After re-training both CNNs and readjusting few parameters (including the focus size range, thresholding step of MSER, and h-minima transform parameter for optimal nucleus separation and focus detection), IRIFs formed by different repair proteins can be analyzed together with their parameters and extent of mutual colocalization in various cell types stained with different methods. On the other hand, the software can be easily tuned to generate comparable results between laboratories for a particular application. This flexibility and robustness, so important especially for research purposes, represents a unique feature of the introduced software.

## Conclusion

Quantification of DNA double-strand breaks by the means of DSB repair focus (IRIF) immunodetection is of utmost importance in various fields of science and practical life (e.g. medicine, cell biology, radiation protection, space exploration, etc.). Because of the nature of IRIFs and fluorescence imaging, where both the IRIF parameters and intensity of analyzed objects may vary dramatically, the automatic segmentation of IRIFs and cell nuclei is highly problematic. We developed a new method based on deep learning that overcomes many of the current limitations of the image analysis and allows rapid and automated quantification and parameter evaluation of IRIF foci. This is enabled by a robust U-Net-based technique for the nucleus segmentation coupled with the U-Net-based focus detection followed by MSER segmentation.

Compared to published approaches, the proposed algorithm works with the 3D confocal multichannel data instead of the single channel 2D slices or maximum image projections. This makes it possible to extract important additional information on morphological and topological IRIF parameters and not only the focus counts. We believe the proposed software, which code is freely available, can substantially simplify the DSB quantification and IRIF analysis. Due to the possibility of the extraction of additional morphometric IRIF and cell nucleus parameters, the software offers numerous practical and research applications. Altogether, the presented software opens the door to a better understanding of IRIF biology and (radiation) DNA damaging and repair.

## Acknowledgements

This work was supported by Czech Science Foundation (projects GACR 20-04109J, GACR 19-09212S), by funds from Specific University Research Grant, as provided by the Ministry of Education, Youth and Sports of the Czech Republic in the year 2020 (MUNI/A/1307/2019 and MUNI/A/1453/2019), by funds from the Faculty of Medicine, Masaryk University to junior researcher (Jaromir Gumulec), 2020, by MEYS CR (Projects 3+3 and Project of Czech Plenipotentiary for cooperation with JINR Dubna) and the Czech-German mobility project DAAD-19-03. We acknowledge the support of NVIDIA Corporation with the donation of the Titan Xp GPU used for this research and OwnCloud storage service provided by CESNET (owncloud.cesnet.cz).

## Data availability

Data used in the manuscript are publicly available in Zenodo repository (www.zenodo.com) and on GitHub (www.github.com). Dataset of Confocal microscopy of gH2AX and 53BP1 DNA repair foci of cells exposed to γ-irradiation – primary cultures and cell lines (DOI 10.5281/zenodo.4067741). Matlab code for automatic segmentation and for labeling, https://github.com/tomasvicar/DeepFoci)

## Abbreviations

53BP1: P53 Binding Protein 1
CNN: Convolutional Neural Network
DSB: DNA double-strand breaks
FOV: Field of View
GUI: Graphical User Interface
IRIF: Ionizing Radiation-Induced (Repair) Foci
MSER: Maximally Stable Extremal Region Algorithm
NHDF: Normal Human Dermal Fibroblasts
RAD51: DNA repair protein RAD51 homolog 1
U-87: U-87 Glioblastoma Cell Line
γH2AX: histone H2AX phosphorylated at serine 139

